## Supplemental Data 1 for "Steroidogenesis and androgen/estrogen signaling pathways are altered in *in vitro* matured testicular tissues of prepubertal mice"

**Supplementary File 1a: Detailed list of antibodies used in this study**

| Primary antibody | Manufacturer | Reference | Dilution | | |
| --- | --- | --- | --- | --- | --- |
|  |  |  | **IF** | **IHC** | **WB** |
| 3β-HSD | Santa Cruz Biot. | sc-515120 (AF488) | 1:100  O/N 4°C |  | 1:1000 5% BSA  O/N 4°C |
| β-Actin | Abcam | ab8226 |  |  | 1:5000 5% BSA  O/N 4°C |
| AR | Abcam | ab133273 | 1:100  O/N 4°C |  | 1:5000 5% milk  O/N 4°C |
| CC3 | Abcam | ab49822 | 1:200  O/N 4°C |  |  |
| CYP17A1 | Abcam | ab231794 | 1:200  O/N 4°C |  | 1:10000 5% milk  O/N 4°C |
| CYP19A1 | BioRad | MCA2077S |  | 1:50  O/N 4°C | 1:250 5% BSA  O/N 4°C |
| FAAH | Proteintech | 17909-1-AP |  |  | 1:1000 5% BSA+5% milk  O/N 4°C |
| Ki67 | Abcam | ab16667 | 1:100  O/N 4°C |  |  |
| **Secondary antibody** | **Manufacturer** | **Reference** |  |  | **WB** |
|  |  |  | **IF** |  |  |
| Goat anti-mouse Alexa 488 | Abcam | ab150013 | 1:200  1h RT |  |  |
| Goat anti-rabbit Alexa 488 | Abcam | ab150077 | 1:200  1h RT |  |  |
| Goat anti-rabbit Alexa 594 | Abcam | ab150080 | 1:200  1h RT |  |  |
| Goat anti-mouse HRP | Invitrogen | 31430 |  |  | 1:5000  O/N 4°C |
| Goat anti-rabbit HRP | Invitrogen | A16110 |  |  | 1:5000  O/N 4°C |

HRP: horseradish peroxidase; IF: immunofluorescence; IHC: immunohistochemistry; O/N: overnight; RT: room temperature; WB: Western Blot

**Supplementary File 1b: List of PCR primers used in this study**

| Gene | Primers sequence | Accession number | Product size (base pair) |
| --- | --- | --- | --- |
| *Abp* | F: GAACGGGGATTCACTGCTG  R: GCAGGCACGAGCGGAAA | NM_011367.2 | 159 |
| *Actb* | F: AGGTCATCACTATTGGCAACGA  R: CACAGGATTCCATACCCAAGAAG | NM_007393.5 | 81 |
| *Ar* | F: CCCGTCCTCTCTGTCTCTGTATAAAT  R: ACAGAGCCAGCGGAAAGTTG | NM_013476.4 | 92 |
| *Cyp11a1* | F: CAGACTTCTTTCGACTCCTCAGAAC  R: CTGGGTGTACTCATCAGCTTTATTGA | NM_001346787.1 | 92 |
| *Cyp17a1* | F: TGCACCGGAAGCTGGTATTC  R: CTGACATATCATCTTCTCCAGTTTCTG | NM_007809.3 | 74 |
| *Cyp19* | F: CCCCTGGACGAAAGTGCTATT  R: CGTTCATACTTTCTGTAGAGCCAAGA | NM_001348171.1 | 119 |
| *Dhh* | F: TGACAGAGCGTTGCAAAGAG  R: GCGCCAACAAACCATACTTA | NM_007857.5 | 195 |
| *Eppin* | F: GGGACCTAGTCTAGCTGACTTGCT  R: GTGTTCACACTCCTCTCTGAATCTG | NM_029325.2 | 67 |
| *Esr1* | F: AAGTGTTACGAAGTGGGCATGA  R: TCTCTGACGCTTGTGCTTCAAC | NM_007956.5 | 84 |
| *Esr2* | F: GGGTGATTTCGAAGAGTGGAATC  R: TTGTTACTGATGTGCCTGACATGA | NM_207707.1 | 97 |
| *Faah* | F: CAGATGGAACACTACAAAGGCTACTT  R: ATACCCTTTTTCATGCCCTTCTTC | NM_010173.4 | 74 |
| *Gapdh* | F: CATGGCCTTCCGTGTTCCTA  R: CCTGCTTCACCACCTTCTTGA | NM_008084.3 | 104 |
| *Gper1* | F: GGGCTAAGCTATGTCATACTCTCAAA  R: AGAGATGACCCCGCAATGTG | NM_029771.3 | 69 |
| *Hsd3b1* | F: AGCATCCAGACACTCTCATCTGACT  R: GTGGGAGCTGGTATGATATAGGGTAA | NM_008293.4 | 79 |
| *Hsd17b2* | F: GGCCGTGGTTAACAATGCC  R: TGAGTTCCCCGTCGATAGGTA | NM_008290.2 | 53 |
| *Hsd17b3* | F: CACTGCAACATTACCTCCGTAGTC  R: ATGAGGCCTTTCCTCCTTGAC | NM_008291.3 | 77 |
| *Igf1* | F: TGGATGCTCTTCAGTTCGTG | NM_001314010.1 | 113 |
|  | R: GCAACACTCATCCACAATGC |  |  |
| *Insl3* | F: TGCTCCTGGCTCTGGGGTCC  R: CACTGCAGCAGCTCCCGGTC | NM_013564.7 | 169 |
| *Rhox5* | F: ATCTCCCTGCACAGTCCTTCA  R: GCTTCCATACCTGGATGATTCTTT | NM_008818.2 | 82 |
| *Septin12* | F: CAACTGAACCCAACCTGGATGTA  R: CCTCGATGACATGGGTCACA | NM_027669.3 | 72 |
| *Srd5a1* | F: GTAACCCATCCCTGTTTCCTG  R: CAGTGCAAAGCCACACCACT | NM_175283.3 | 198 |
| *Star* | F: AGCAAAATGACTCAGGGTGTAAGTG  R: TTCATCTCTCTCCTTCCATCTCTGT | NM_011485.5 | 81 |
| *Sult1e1* | F: TGTTGAAATGTTCTTGGCAAGGCC  R: CATCCTCCTTGCATTTTTCCACATCA | NM_023135.2 | 131 |

**Supplementary File 1c: Compound-specific MRM parameters for LC-MS/MS**

| **Name Q = Quantifier / q = qualifier** | **Precursor ion (m/z)** | **Product ion (m/z)** | **Q1 Pre Bias (V)** | **CE** | **Q3 Pre Bias (V)** |
| --- | --- | --- | --- | --- | --- |
| Androstenedione-13C3-Q | 290.1 | 100.15 | -10 | -22 | -19 |
| Androstenedione-13C3-q | 290.1 | 112.2 | -17 | -23 | -12 |
| Androstenedione-Q | 287.1 | 97.15 | -16 | -22 | -19 |
| Androstenedione-q | 287.1 | 109.2 | -17 | -24 | -21 |
| DHEA-D5-Q | 276.1 | 218.2 | -17 | -19 | -16 |
| DHEA-D5-q | 276.1 | 202.25 | -17 | -22 | -14 |
| DHEA-Q | 271.1 | 213.3 | -17 | -17 | -16 |
| DHEA-q | 271.1 | 197.25 | -16 | -20 | -14 |
| Testosterone-13C3-Q | 292.1 | 112.1 | -18 | -23 | -23 |
| Testosterone-13C3-q | 292.1 | 100.15 | -18 | -23 | -11 |
| Testosterone-Q | 289.1 | 109.2 | -17 | -26 | -12 |
| Testosterone-q | 289.1 | 97.15 | -17 | -22 | -18 |

CE: collision energy; DHEA: Dehydroepiandrosterone; MRM: multiple reaction monitoring.
